# Role for both myeloid and plasmacytoid DC in driving primary T helper cell response to *Burkholderia pseudomallei* in healthy individuals

**DOI:** 10.1101/2020.06.22.164608

**Authors:** Durga Reddi, Lydia Durant, David Bernardo, Alistair Noble, Nicholas R. English, Philip Hendy, Graeme C. Clark, Joann L. Prior, E. Diane Williamson, Stella C. Knight

## Abstract

*Burkholderia pseudomallei* is a Gram-negative bacterium that causes melioidosis, an infectious disease endemic to south-east Asia. As *B. pseudomallei* is antibiotic-resistant, the need for cell-based vaccines and therapies is crucial to managing melioidosis. Dendritic cells (DC) provide the first line of defense to infection and direct downstream immune responses. Using practical volumes of fresh healthy donor blood, we show that heat-killed *B. pseudomallei* activated and stimulated expression of pro-inflammatory cytokines TNF-α, IL-1β and IL-6 from both myeloid and plasmacytoid DC. Furthermore, *B. pseudomallei-*pulsed DC induced activation and proliferation of CD4^+^ T-cells. Thus, both DC subsets are important for driving primary T helper cell responses to *B. pseudomallei* in healthy individuals and have the potential to be targeted for future therapies and vaccines.

**Author Summary:** Melioidosis is an infectious disease endemic to south-east Asia and northern Australia caused by the bacterium *Burkholderia pseudomallei*. Melioidosis presents a significant public health threat because it has no effective vaccine or cure, leading to a high mortality rate of between 10-50%. We highlight the possibility of immune-based strategies targeting *Burkholderia pseudomallei* to better treat and prevent melioidosis. Specifically, we show that dendritic cells-the sentinel cells of the immune system-respond to *B. pseudomallei* in healthy individuals and in turn can orchestrate downstream protective immune responses. Thus, dendritic cells may be key players in the development of both vaccines and therapeutics for melioidosis as well as other bacteria-driven diseases.

## Introduction

*Burkholderia pseudomallei* (B.psm) is a Gram-negative infectious bacterium and the causative agent of melioidosis, a disease endemic to parts of south-east Asia. Disease may present as a chronic or acute infection of skin, lung, liver or spleen and can lead to pneumonia and septicemia (1). After initial exposure, B.psm replicates intracellularly within epithelial or phagocytic cells and can evade host immunity (2). Due to the multiple routes of exposure, active efflux of antibiotics and lack of an available vaccine, infection with B.psm is problematic to manage and in need of improved post-exposure therapy.

Dendritic cells (DC) are the first line of defense to infection and provide a link between innate and adaptive immunity. DC express pattern recognition receptors (PRR) which recognise pathogen-associated molecular patterns (PAMPs) within microbes. DC internalise microbial components by receptor-mediated endocytosis, then process and present antigens associated with major histocompatibility complex (MHC) class II to CD4^+^ or MHC class I to CD8^+^ T-cells residing in local lymph-nodes. Upon stimulation, DC also express CCR7, allowing their migration to local lymph-nodes, and maturation markers including CD80, CD86 and CD40 which provide co-stimulatory signals to T-cells. Finally, DC secrete a variety of cytokines which direct specific T-cell responses.

The two major subsets of DC in humans are myeloid and plasmacytoid. Myeloid DC (mDC) are typically responsive to bacteria whereas plasmacytoid DC (pDC) respond to viruses (3), although pDC can be activated by some bacteria including *Staphylococcus aureus* and *Streptococcus pyogenes* (4)(5). Moreover, pDC co-cultured with mDC responsive to bacterial stimulation show a unique capacity to mature and be activated, suggesting synergy between the two subsets (6). Human and murine pDC internalise and cause killing of B.psm (7). Furthermore, vaccination of mice with activated bone marrow (BM) DC could induce protective immunity against subsequent infections with B.psm, demonstrating the potential for DC-based vaccines (8).

The focus of this study was to investigate primary human DC responses to B.psm to contribute towards developing effective cellular immunotherapies. Total DC were enriched from fresh PBMC and pulsed with heat-killed B.psm to assess contributions of both subsets in priming immune responses by examining their phenotype (activation and cytokine expression) and function (ability to stimulate primary T-cell responses). We highlight roles for both mDC and pDC in stimulating primary T cell responses to B.psm. Furthermore, our methodology can be used to assess memory responses in exposed individuals or to screen for bacterial sub-components to be used in vaccines.

## Results

### *B. pseudomallei* activates both mDC and pDC enriched from fresh blood

Immunisation with heat-killed B.psm protects mice against subsequent pathogen challenge, suggesting that killing of the bacteria preserves protective motifs and antigens (9). To define an optimal non-toxic concentration of B.psm, DC enriched from fresh PBMC were incubated with ascending concentrations of B.psm for 20 h, with a baseline 0 h time-point used for comparison. Expression of activation markers CD80, CD86 and CD40 and lymph node migration marker CCR7 were measured on mDC and pDC by flow cytometry. The proportion of mDC expressing CD80, CD86 and CD40 was significantly increased in response to high concentrations of B.psm at 20h (Fig 1A). The highest concentration of B.psm also stimulated a significant increase in the proportion of pDC expressing CD80, but not CD86 or CD40. (Fig 1B). Interestingly, the proportion of mDC expressing CCR7 increased in all groups at 20h, the proportion of pDC expressing CCR7 was lower at 20h in all groups; these changes were not statistically significant (Supplementary Fig 3). However, median fluorescence intensity (MFI) of HLA-DR (MHCII) on both mDC and pDC increased significantly after 20h of B.psm incubation (Fig 1C). Thus, we determined that the highest concentration of 1 × 10^6^ CFU/ml of B.psm optimally activated both DC subsets and was used in subsequent experiments.

**Fig 1.**
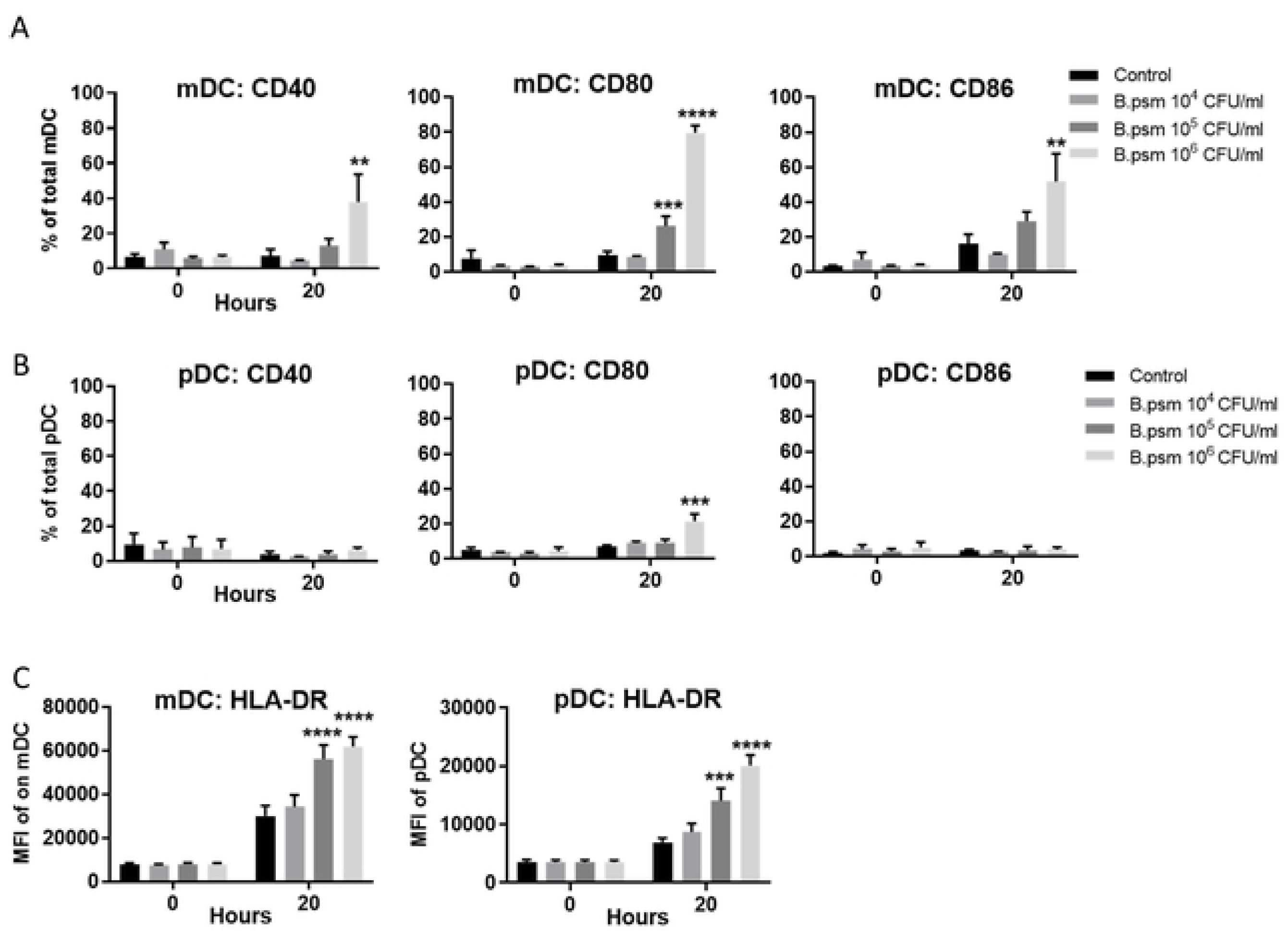
*B. pseudomallei* stimulates maturation and cytokine production by myeloid and plasmacytoid DCs in healthy donors. Enriched DC from healthy donor PBMC were stimulated 20 h with *B. pseudomallei* (B.psm) and percentage of cells expressing CD80, CD86 and CD40 was measured for **(A)** myeloid (m)DC and **(B)** plasmacytoid (p)DC by flow cytometry. **(C)** The median fluorescence intensity (MFI) of HLA-DR on mDC and pDC was also determined at 20h. Bars represent mean values ± SEM from three (A-B) or six (C) independent experiments. A two-way (ANOVA) followed by Dunnett’s multiple comparisons test was used to compare experimental groups to the medium only control. ** *p* <0.01, *** *p* <0.001, **** *p* <0.0001.

### B.psm stimulates proinflammatory DC cytokine production

Activated DC secrete a variety of polarising cytokines that direct protective immune responses. We investigated production of TNF-α, IL-1β and IL-6 by DC in response to B.psm. At baseline, there was no detectable TNF-α, IL-1β or IL-6 in supernatants of stimulated DC (Fig 2A). However, after 24 h of pulsing with B.psm, TNF-α and IL-6 were significantly increased (Fig 2A). To determine which DC subset was synthesising these cytokines, we stimulated DC for 3h with B.psm and measured intracellular cytokines in mDC and pDC. B.psm stimulation led to a significant increase in proportion of mDC expressing TNF-α and IL-1β, while mDC expressing IL-6 was only significantly increased by *E. coli* LPS (Fig 2B). B.psm also stimulated a significant increase in proportion of pDC expressing TNF-α.(Fig 2C). The proportion of mDC or pDC expressing TNF-α was similar (approximately 20%), but MFI of TNF-α was significantly higher in mDC compared to pDC in response to both B.psm (1.78 fold-higher) and *E. coli* LPS (1.46 fold-higher) stimulation (Supplementary Fig 4).

**Fig 2.**
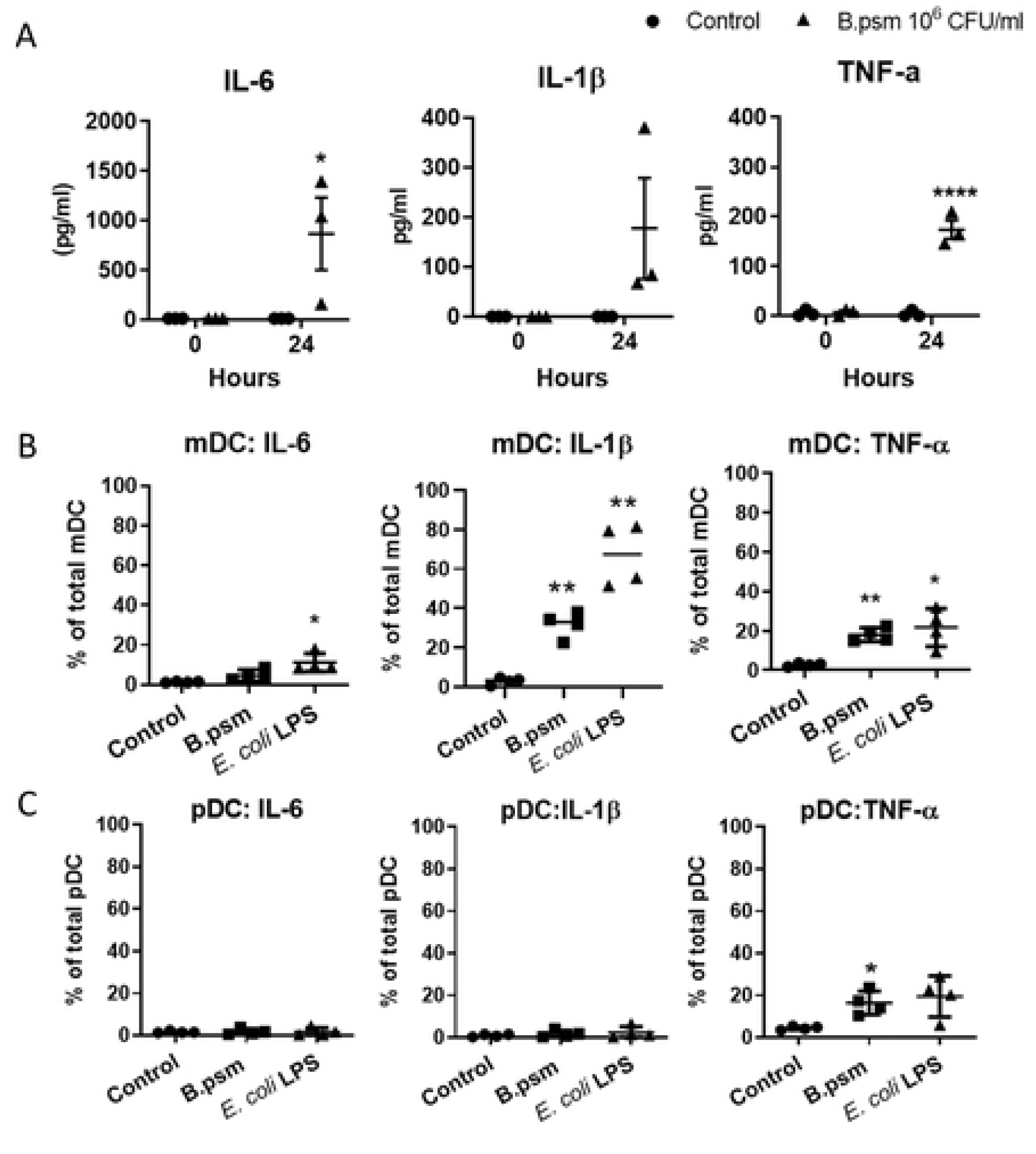
B. pseudomallei induces inflammatory cytokine production by myeloid and plasmacytoid DC in healthy donors. (**A**) Amounts of secreted TNF-α, IL-1β and IL-6 in supernatants of enriched DC following 24 h stimulation with B.psm or medium only are shown. Percentage of (**B**) mDC and **(C)** pDC expressing intracellular IL-6, IL-1β and TNF-α following 3h stimulation with B.psm, *E. coli* LPS or medium only. Bars represent mean values ± SEM from three (A) or four (B-C) independent experiments. A one-way (ANOVA) followed by a Dunnett’s multiple comparisons test was used to compare experimental groups to the medium only control. * *p*< 0.05, ** *p* <0.01, **** *p* <0.0001.

### DC conditioned with B.psm stimulate CD4^+^ T-cell proliferation

B.psm-stimulated DC became activated and produced cytokines, so we next assessed their ability to induce T-cell proliferation. Enriched DC were cultured with B.psm for 20h and added at 5% or 10% to syngeneic CD4^+^ and CD8^+^ T-cells for 7 days. T-cells cultured with anti-CD3 and anti-CD28 were used as a positive control. Divided T-cells were defined as CD4^+^ or CD8^+^ and CTV^low^ (Fig 3A). T-cell markers, CD45RA and CD45RO were used to confirm naïve or activated/memory status, respectively, within the divided and undivided cells (Fig 3A). B.psm-conditioned DC added at 5% induced CD4^+^ T-cell proliferation which was significantly higher than the untreated control after 7 days culture (Fig 3B). Conversely, B.psm*-*pulsed DC added at either 5% or 10% did not significantly induce CD8^+^ T-cell proliferation (Fig 3C). The majority of CD4^+^ T-cells that proliferated in response to B.psm treatment were CD45RO^+^, indicating activation of these cells (Fig 3A). Thus, primary response to B.psm in unexposed individuals involves a CD4^+^ T helper cell response, with less indication for CD8^+^ T-cell involvement.

**Fig 3.**
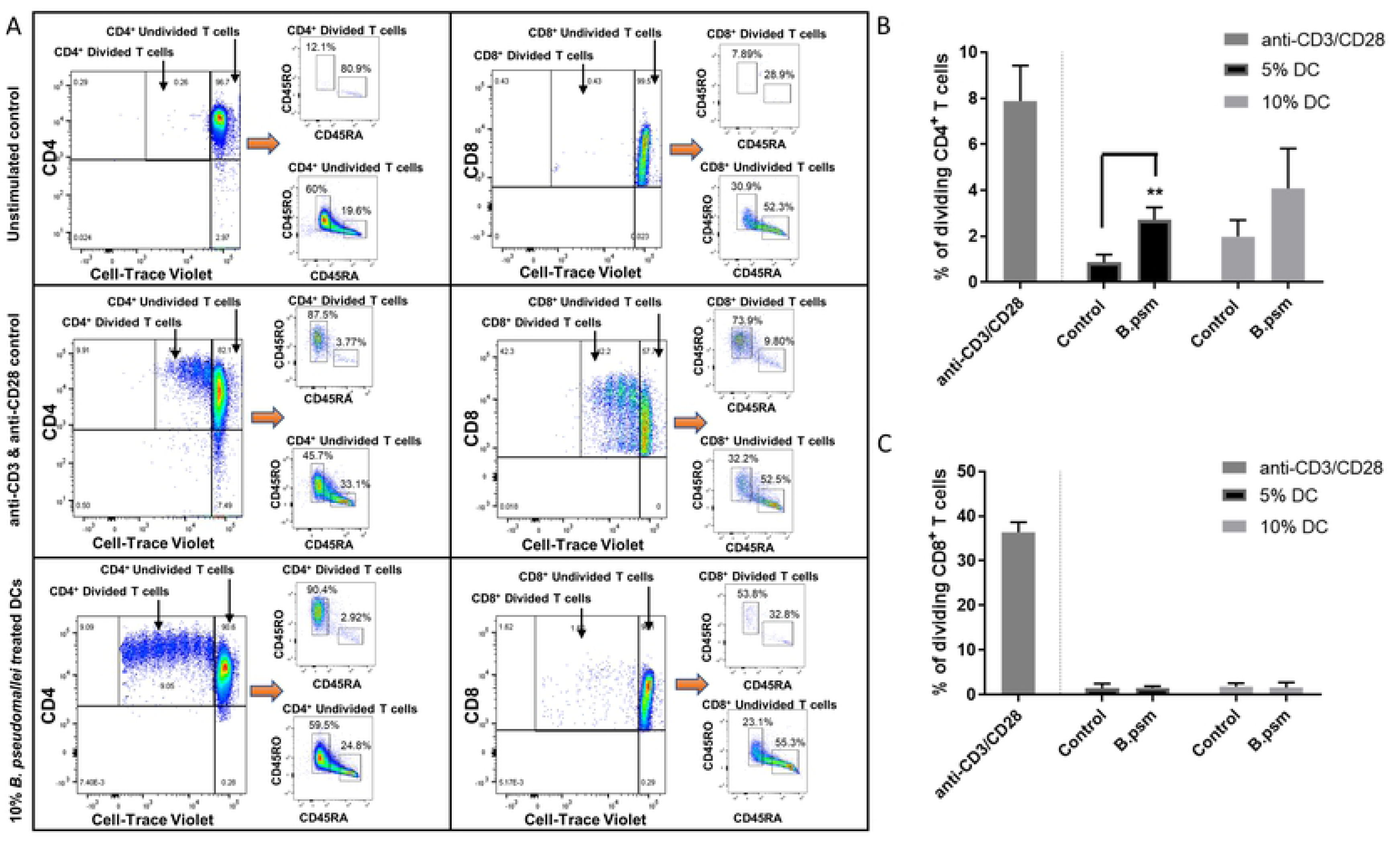
*B. pseudomallei* stimulates a primary CD4^+^ T helper cell response in healthy donors. Enriched DC were stimulated for 20h with B.psm or medium alone (control). 2.5 x 10^5^ syngeneic total T-cells were then co-cultured with either 5% or 10% stimulated DC for 7 days. Anti-CD3 and anti-CD28 were used as a positive control for T-cell stimulation. **(A)** Gating on Cell-Trace Violet^low^ CD4^+^ (left panels) or CD8^+^ (right panels) cells defines the divided T-cell populations. Examples are given of unstimulated (top panels), positive stimulation with anti-CD3 & anti-CD28 (middle panels) or 10% DC-treated with B.psm (bottom panels) CD4^+^ and CD8^+^ T-cells after 7 days culture. Expression of cell-surface markers CD45RO and CD45RA were used to define activation status of proliferated vs. non-proliferated T-cells. Pooled data showing the % of proliferated **(B)** CD4^+^ or **(C)** CD8^+^ T-cells after 7 days incubation with 5% or 10% of DC stimulated with or without B.psm or with anti-CD3 and anti-CD28. Bars represent mean values ± SEM from three independent experiments. A Student’s paired T*-*test was used to compare the untreated control to B.psm group. ** *p* <0.01.

## Discussion

We demonstrate that heat-killed B.psm activates both mDC and pDC enriched from fresh PBMC of healthy individuals and induces their production of pro-inflammatory cytokines TNF-α and IL-1β. While mDC were more strongly activated than pDC, with greater expression of costimulatory molecules and IL-1β, both subsets showed a significant up-regulation of MHC-II and produced TNF-α in response to B.psm. Furthermore, total mDC and pDC stimulated with heat-killed B.psm stimulated proliferation of syngeneic CD4^+^ T-cells *in vitro*, suggesting that downstream immunity involves a protective T helper cell response.

Our study used modest volumes of blood from single healthy donors to examine immune response to B.psm, which sets it apart from previous studies using bulk buffy coats [9]. We chose to enrich DC from PBMC using a negative selection protocol because, although positive selection gave higher purity, negatively enriched DC survived longer in culture. Conveniently, use of negatively-selected DC allowed us to study both DC subsets simultaneously and build on previous work suggesting that pDC as well as mDC are directly involved in the immune response to B.psm (7). Interestingly, both CD4^+^ and CD8^+^ T-cell responses were induced when whole PBMC from recovered melioidosis patients were cultured with B.psm or purified proteins, suggesting that protective memory involves both T-cell types (10). It will be important in the future to dissect further the specific functions of mDC and pDC in promoting protective immune responses to B.psm to develop more effective vaccines and therapies.

DC-based vaccines have been used to treat cancer and autoimmune conditions (11). More recently, pDC were used to treat patients with metastatic melanoma by loading them with tumour antigens to induce type I interferon expression (12). Due to the low numbers of DC present in peripheral blood, *ex vivo* loading of DC with antigen for vaccine administration is not feasible in terms of cost, labour and the need for tailored therapy for each patient. Targeting of antigen to DC *in vivo* using antigen-antibody complexes to DC cell-surface receptors such as CD205 may prove feasible (13). Live B.psm are taken up by DC and induce maturation and migration of immature DCs *in vitro* (14). Although the interaction with the host cell differs for live and killed bacteria, the ability of DC to become activated by heat-killed B.psm and our demonstration that these activated DC can in turn activate even unprimed T-cells, suggests that DC-based vaccines could be efficacious in treating or preventing B.psm infection.

By using cells from individuals with suspected melioidosis, it will be possible to apply methods described herein to test for memory responses not only to killed bacteria, but also to bacterial sub-components. Using cells from convalescent patients may identify components required for protective immune responses. Matching requirements of protective immunity to the ability of antigenic bacterial components to produce primary, pro-inflammatory activation in DC, will facilitate the identification of effective vaccine sub-units and enable development of improved prophylactic strategies.

## Materials and Methods

### Bacterial preparation

*B. pseudomallei* K96243 was cultured in L-broth and on L-agar at 37°C. Bacterial stocks were maintained at –80°C as 20% glycerol suspensions. *B. pseudomallei* K96243 was handled at Advisory Committee for Dangerous Pathogens (ACDP) containment level 3. The bacteria were heat-killed as previously reported (15).

### Healthy Donors

Blood (30ml-50ml) was collected from healthy volunteers with no known autoimmune or inflammatory diseases, allergies, or malignancies. Ethical approval was obtained from the Health Research Authority UK and London Brent Research Ethics Committee (05.Q0405.71). Written informed consent was received from participants prior to inclusion.

### Dendritic cell enrichment

Peripheral blood mononuclear cells (PBMCs) were isolated by centrifugation over Ficoll-Paque Plus (Amersham Biosciences, Chalfont St. Giles, UK). Total mDC and pDC were negatively enriched from PBMCs using the EasySep™ Human Pan-DC Pre-Enrichment Kit (StemCell Technologies) according to manufacturer’s instructions. Compared to whole PBMC where DC (HLA-DR^+^ and lineage (CD3/CD14/CD16/CD19/CD34)^-^ cells) are between 1-2% of live cells, the purity of DC (mDC and pDC) after enrichment was between 65-70% of live cells (Supplementary Fig 1A and B). Enriched DC were cultured in Dutch modified RPMI 1640 (Sigma-Aldrich, Dorset, UK) containing 100 U/mL penicillin/streptomycin, 2 mM L-glutamine, 50 μg/mL gentamicin (Sigma-Aldrich) and 10% fetal calf serum (TCS cell works, Buckingham, UK).

### Bacterial stimulation

Enriched DC (5 × 10^4^) were stimulated with ascending concentrations of B.psm or medium only (untreated control) for 0h, 3h, 20h or 24h at 37 °C and 5% CO_2._ For DC intracellular cytokine analysis, stimulation with 100ng/ml of *E. coli* LPS (Sigma-Aldrich, Dorset, UK) was a positive control.

### Cell-surface antibody labelling

After 20h incubation, DC were washed with FACS buffer (1x PBS containing 5% FCS, 1mM EDTA and 0.02% sodium azide) and labelled with Near-IR Live/Dead Fixable Dead Cell stain (ThermoFischer Scientific) according to kit instructions. Cells were then labelled for 20 minutes on ice with antibodies (Table S1), fixed with 1% paraformaldehyde in 0.85% saline and stored at 4°C until flow cytometric analysis. “Fluorescence minus one” controls were used to determine positive staining for each marker.

### Intracellular cytokine analysis

After 3h culture, DC were harvested, labelled with viability stain and surface antibodies as above and fixed and permeabilised using Leucoperm A and B reagents (Bio-Rad, Watford, UK). Cells were labelled intracellularly with antibodies (Table S2) in Leucoperm B. Positive staining for all cytokines was determined by comparing to an untreated control.

### Enzyme linked immunosorbent assay (ELISA)

Enriched DC were cultured with B.psm for 24h and supernatants stored at −20°C. Concentrations of IL-6, IL-1β and TNF-α were measured using DuoSet ELISA kits (R&D Systems, Abingdon, UK) as per manufacturers’ instructions.

### DC: T cell co-cultures

Enriched DC were stimulated with B.psm for 20 h. Total T-cells were purified from the same donor’s PBMC using the EasySep™ Human T cell Isolation kit (StemCell Technologies). Purity of total CD4^+^ and CD8^+^ T-cells was ∼94% (Supplementary Fig 2). After 20h culture, T-cells were labelled with Cell Trace Violet (CTV) dye (Thermo Fisher) as per manufacturer’s instructions. CTV-labelled T-cells (2.5 × 10^5^) were co-cultured with either 10% (2.5 × 10^4^) or 5% (1.25 × 10^4^) B.psm-pulsed DC for 7 days. As a positive control, 2.5 × 10^5^ CTV-labelled T-cells were stimulated with 5µg/ml of plate-bound anti-CD3 (300331, Biolegend) and 5µg/ml of soluble anti-CD28 (302913, Biolegend). CTV-labelled T-cells cultured in medium only were an unstimulated control.

### Flow cytometry analysis

Cells were acquired on the FACSCanto II Flow Cytometer (BD Biosciences) and analysed using FlowJo software version 10.

### Statistical Analysis

One-way or two-way ANOVA followed by Dunnett’s multiple comparisons test was used to compare experimental groups to medium only controls. A Student’s paired T-test was used to compare T-cell proliferation in the B.psm group to untreated control. Statistical analysis used GraphPad Prism version 7.0.

## Acknowledgments

We thank Dr. Rakesh Vora for his assistance with recruitment of healthy volunteers.

## Supporting Figure Captions

**S1 Fig. Enrichment of human blood myeloid and plasmacytoid dendritic cells from fresh PBMC.** DC were identified using side/forward scatter properties, doublet discrimination and gating on viable cells. Cells that were HLA-DR^+^ and lineage^−^ (CD3, CD14, CD16, CD19 and CD34) were further divided into mDC (CD11c^+^CD123^-^) and pDC (CD11c^-^CD123^+^). (A) DC identification within PBMC population before DC enrichment. (B) DC identification after enrichment.

**S2 Fig. Purification of Total CD4**^**+**^ **and CD8**^**+**^ **T-cells from fresh PBMC.** Total T-cells were purified on day 0 from PBMC and identified using side/forward scatter properties, doublet discrimination and total viable cells. Total CD3^+^ T-cells were subdivided using CD4 and CD8 markers to distinguish between CD4^+^ and CD8^+^ T-cells respectively.

**S3 Fig. Lymph-node homing marker CCR7 is induced nonspecifically by *B. pseudomallei.*** Enriched blood mDC and pDC from healthy donors were stimulated with ascending concentrations *B. pseudomallei* (B.psm) for 20 h and expression of lymph node homing marker CCR7 on DC was assessed by flow cytometry. Graphs show the frequency of (A) mDC and (B) pDC expressing CCR7 after B.psm stimulation. Bars represent mean values ± SEM from three independent experiments.

**S4 Fig. TNF-α is more highly upregulated in mDC than pDC in response to *B. pseudomallei*.** MFI of TNF-α in mDC vs. pDC after 3h stimulation with untreated control, 10^6^ CFU/ml B.psm or 100 ng/ml *E. coli* LPS. Bars represent mean values ± SEM from four independent experiments. A one-way (ANOVA) followed by a Dunnett’s multiple comparisons test was used to compare experimental groups to the media only control. * *p*< 0.05, ** *p* <0.01.

